# Circadian clock neurons mediate time-of-day dependent responses to light

**DOI:** 10.1101/291955

**Authors:** Jeff R. Jones, Tatiana Simon, Lorenzo Lones, Erik D. Herzog

## Abstract

Circadian (~24 h) rhythms influence nearly all aspects of physiology, including sleep/wake, metabolism, and hormone release. The suprachiasmatic nucleus (SCN) synchronizes these daily rhythms to the external light cycle, but the mechanisms by which this occurs is unclear. The neuropeptide vasoactive intestinal peptide (VIP) is the predominant contributor to synchrony within the SCN and is important for circadian light responses, but the role of VIP neurons themselves is unclear. Thus, we tested the hypothesis that rhythmic SCN VIP neurons mediate circadian light responses. Using *in vivo* fiber photometry recording of SCN VIP neurons we found daily rhythms in spontaneous calcium events that peaked during the subjective day and in light-evoked calcium events that exhibited the greatest response around subjective dusk. These rhythms were correlated with spontaneous and NMDA-evoked VIP release from SCN VIP neurons *in vitro*. Finally, *in vivo* hyperpolarization of VIP neurons attenuated light-induced shifts of daily rhythms in locomotion. We conclude that SCN VIP neurons are circadian and depolarize to light to modulate entrainment of daily rhythms in the SCN and behavior.

## INTRODUCTION

Organisms have evolved endogenous circadian (~24 h) rhythms in behavior and physiology, including sleep/wake, hormone release, and metabolism (Kalsbeek et al. 2006), to anticipate reliable daily events such as the light cycle. In mammals, light at night but not during the day shifts a master circadian pacemaker in the suprachiasmatic nucleus (SCN) to entrain daily rhythms to local time (Gillette, 1996; Sakamoto et al., 2013). Chronic nocturnal light exposure is associated with an increased risk of disease (Chang et al., 2015; Fonken et al., 2010; LeGates et al., 2012) and disruption of circadian rhythms in the SCN and behavior (Granados-Fuentes et al., 2004; Ohta et al., 2005). The mechanisms of normal and pathological phototransduction in the SCN, however, are unclear. The neuropeptide vasoactive intestinal peptide (VIP), produced by retinorecipient neurons comprising ~10% of the SCN network, plays a critical role in synchronizing SCN cells to each other and to the the light cycle (Abrahamson and Moore, 2001; Aton et al., 2005; Colwell et al., 2003; Maywood et al., 2011). For instance, VIP administration shifts daily rhythms both *in vitro* and *in vivo* (An et al., 2011; Atkins et al., 2010; Piggins et al., 1995; Reed et al., 2001), and *Vip*^*-/-*^ mice show deficits in their circadian responses to light, such as entraining to light with an 8 h advanced phase (Colwell et al., 2003; Vosko et al., 2015). Here we tested the hypothesis that SCN VIP neurons regulate light responses with time of day to promote shifts in daily behavior. We found that SCN VIP neurons are circadian and excited by light *in vivo,* release VIP depending on time of day and excitatory input, and are necessary for the normal transduction of light signals to the SCN at times when light shifts daily rhythms.

## RESULTS

To determine the circadian regulation of SCN VIP neuron activity over multiple days, we used long-term *in vivo* fiber photometry to record hourly spontaneous calcium activity from the SCN of freely-moving mice. We injected a Cre-dependent adeno-associated virus encoding the fluorescent calcium sensor GCaMP6s (AAV9-CAG-Flex-GCaMP6s) or a control fluorophore (AAV9-CAG-Flex-EGFP) into the SCN of VIP-IRES-Cre knock-in mice (**Figs. 1a, b; Supplementary Figs. 1a-d**). We found daily rhythms in intracellular calcium levels and event frequency that persisted for as many days as we recorded, peaking during the daily minimum of locomotor activity in a light-dark (LD) cycle or in constant darkness (DD; **Fig. 1c-d; Supplementary Figs. 1e-f**). The amplitude of the rhythm (LD day, night = 1.31 ± 0.38, 0.09 ± 0.15 events/min; DD subjective day, night = 1.34 ± 0.48, 0.09 ± 0.15 events/min) was greatly disrupted in constant light (LL day, night = 1.34 ± 0.53, 1.41 ± 0.58 events/min). We therefore tested how SCN VIP neurons responded to light and found that a 10-min light pulse reliably increased intracellular calcium (**Supplementary Fig. 1g**). We next tested the effects of 15-s light pulses at different times of day and found that calcium responses exceeded baseline during the subjective night (circadian times, CT, 12, 18 or 24), but not during the subjective day (CT 6; **Fig. 1e; Supplementary Fig. 1h**). These responses lasted multiple tens of seconds after the cessation of the light pulse, with daily peaks in amplitude and duration occurring around CT 12 (**Figs. 1f, g; Supplementary Video 1**) and increased in amplitude with increasing light intensity (**Supplementary Figs. 1i,j**). These results suggest that SCN VIP neurons exhibit *in vivo* daily rhythms in spontaneous and light-evoked calcium activity and can encode light intensity in a manner consistent with light-induced circadian phase shifts (Nelson and Takahashi, 1991).

**Figure 1.**
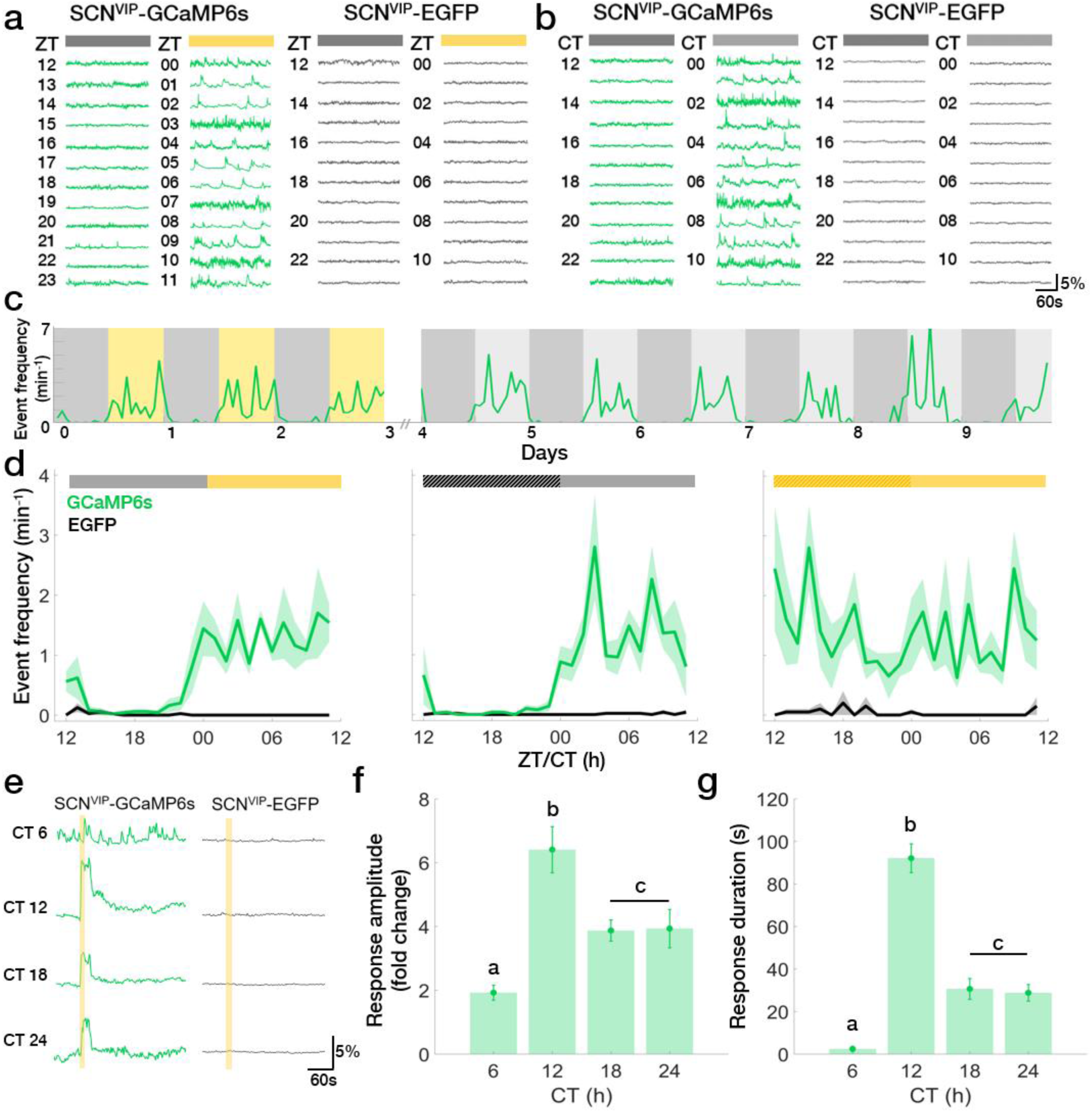
SCN VIP neurons exhibit daily rhythms in spontaneous and light-evoked calcium activity *in vivo*. **a-b**) Hourly ΔF/F traces taken from VIP neurons in transgenic mice with (GCaMP6s) or without (EGFP) a fluorescent calcium reporter in a daily light-dark cycle (LD; yellow = light phase) or constant darkness (DD; light grey = subjective day). ZT, zeitgeber time; CT, circadian time. **c**) Long-term recordings revealed reliable daily rhythms in the rate of VIP neuron calcium events from a representative GCaMP6s^+^ mouse in LD and DD. **d**) The average daily profile of VIP neuron calcium event frequencies depended on circadian time and presence of light in LD, DD, or constant light (LL; light yellow = subjective day) in GCaMP6s (green; n = 5, 5, 4 mice; JTK Cycle, p<0.001), but not EGFP (black; n = 5, 4, 2; JTK Cycle, p>0.05) mice. Light during subjective night in LL increased event frequency compared to DD or LD (Two Way ANOVA, F(2, 330) = 19.48, p<0.001 with post-hoc Tukey’s MCT). Lines and shadings depict mean ± SEM. **e**) Representative calcium responses to a 15-s light pulse presented to representative GCaMP6s and EGFP mice at different circadian times. **f-g**) The amplitude and duration of the light-evoked calcium response of VIP neurons at different CTs peaked around subjective dusk (n = 4 mice, 2 trials per CT; One-Way Repeated Measures ANOVA, F(3, 21) = 14.15, p < 0.0001; F(3, 21) = 67.08, p<0.0001 with post-hoc Tukey’s MCT). Error bars depict means ± SEM; letters indicate statistically different groups.

Next, to determine how SCN VIP neuron calcium activity correlates with the release of VIP, we engineered HEK293 reporter cells to selectively fluoresce in response to VIP. These cells reported VIP concentration in a dose-dependent manner (**Supplementary Fig. 2a-c**) and exhibited spontaneous fluorescence when cultured under acute SCN slices, but not when in the same dish remote from the SCN (**Supplementary Fig. 2d, e**). VIP sensors exhibited more spontaneous calcium activity when placed under acute wild-type SCN slices during the subjective day than during the subjective night (**Fig. 2a**). Spontaneous calcium activity was circadian when recorded under cultured wild-type SCN slices, but was arrhythmic and low amplitude when recorded under *Vip*^-/-^ slices (**Fig. 2b; Supplementary Fig. 2f; Supplementary Video 2**). Adding 1 or 10 mM NMDA to acute wild-type SCN slices at CT 12 to mimic light input (Ding et al., 1994; Mintz et al., 1999) also increased calcium activity in VIP sensors (**Fig. 2d, e, Supplementary Video 3**). Consistent with the inability of daytime light to shift the SCN, adding 1 mM NMDA to the SCN at CT 6 did not change VIP sensor fluorescence. Importantly, adding NMDA to reporter cells alone or to a *Vip*^-/-^ SCN slice did not change spontaneous fluorescence. Intriguingly, 10 mM NMDA reduced VIP sensor fluorescence (which could be restored by adding VIP). These results collectively suggest that there are daily rhythms in spontaneous and NMDA-evoked VIP release from the SCN, consistent with a role for VIP in mediating circadian-, light- and NMDA-induced shifts of the SCN and behavior.

**Figure 2.**
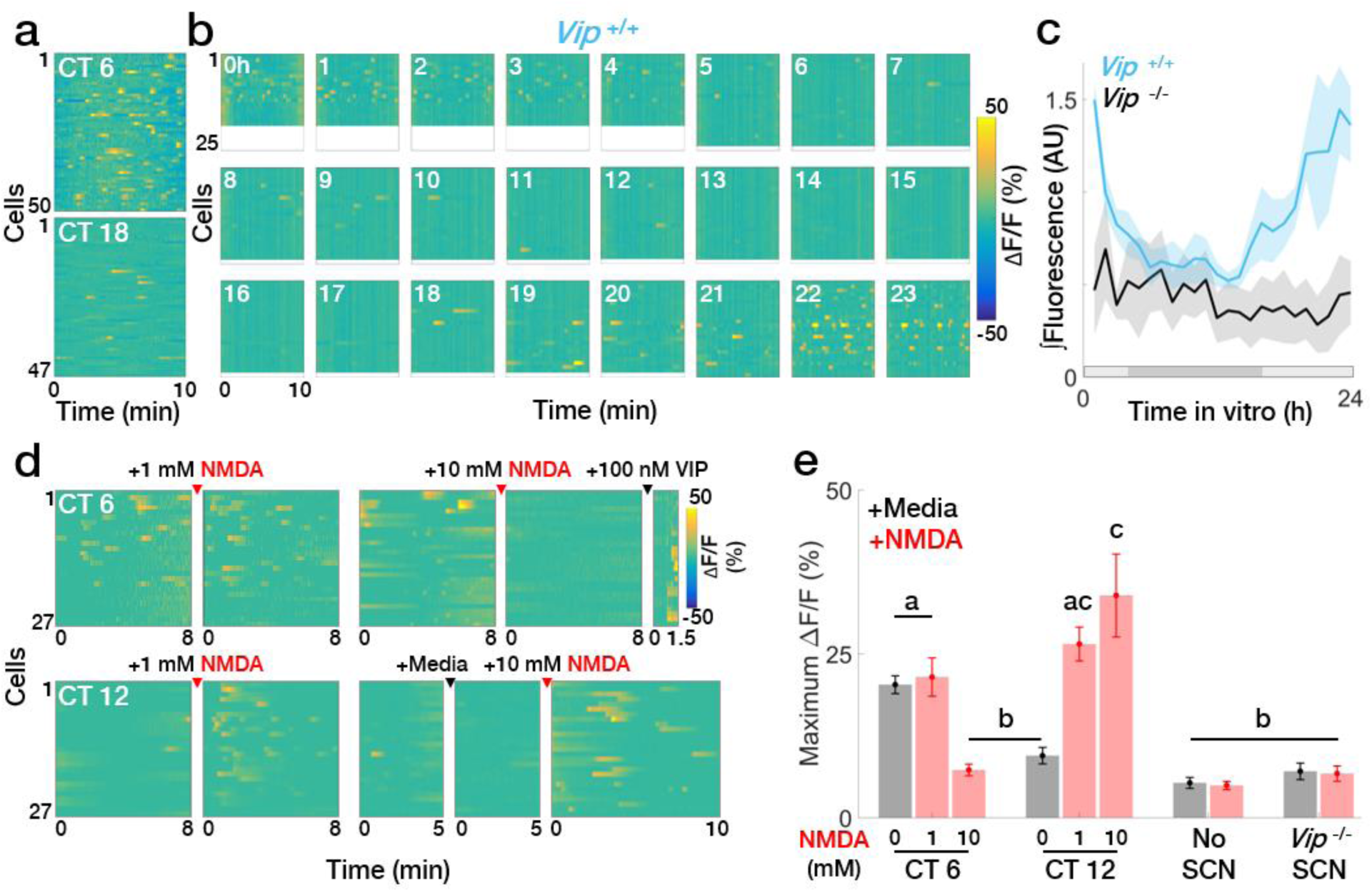
SCN slices exhibit daily rhythms in spontaneous and evoked VIP release. **a**) Representative heatmaps of ΔF/F responses from VIP sensors cultured under acute SCN slices explanted during the subjective day or night. **b-c**) Hourly ΔF/F heatmaps and summary plot showing the daily rhythm in average calcium responses integrated over 10 min from individual reporter cells cultured under an organotypic wild-type SCN slice. Shaded bar depicts the extrapolated subjective day (light gray) and night (dark gray; *Vip*^+/+^ n = 5 slices, JTK Cycle, p<0.001; *Vip*^-/-^ n = 5 slices, JTK Cycle, p>0.05). **d-e**) Heatmaps and summary plots of ΔF/F responses to 1 μl of 1 or 10 mM NMDA applied to acute SCN slices cultured over VIP sensors at CT 12 or 6 (n = >100 cells, 4 dishes per group; Two-Way ANOVA, F(9, 36) = 17.29, p < 0.0001 with post-hoc Tukey’s MCT). Error bars depict means ± SEM; letters indicate statistically different groups.

Finally, to determine the contribution of SCN VIP neurons to the encoding of light that shifts daily rhythms, we used chemogenetics to acutely silence these neurons in freely-behaving mice. We recorded circadian locomotor behavior from VIP-IRES-Cre mice injected with an SCN-targeted, Cre-dependent AAV encoding an engineered inhibitory Gi-coupled receptor (AAV8-hSyn-DIO-hM4Di) or a control fluorophore (AAV9-CAG-Flex-EGFP; **Supplementary Fig. 3a**). We then silenced VIP neurons around the daily time of locomotor onset (CT 12), when *in vivo* VIP neuron calcium activity is decreasing and could be augmented by light exposure. Intraperitoneal injections of the hM4Di-specific ligand clozapine-N-oxide (CNO) at CT 11.5 did not shift daily rhythms in locomotor activity in experimental or control mice (**Fig. 1d**), indicating that VIP neuron activity is not required for circadian rhythms to progress in constant conditions. Strikingly, injection of CNO 30 min prior to a 15-min light pulse at CT 12 greatly attenuated the resulting phase delay from 2.10 ± 0.21 h to 0.88 ± 0.07 h compared to control animals (**Figs. 3a, b**). We also found CNO similarly reduced phase delays of circadian behavior after light pulses that were 10-times dimmer (0.39 ± 0.04 h compared to shifts of CNO-treated control mice 0.91 ± 0.04 h; **Supplementary Figs. 3b, c**) and attenuated light induction of c-FOS, an indicator of neuronal activation, in the SCN (**Figs. 3c, d; Supplementary Fig. 3d**). These results suggest that depolarization of SCN VIP neurons is necessary for normal light-induced phase shifts.

**Figure 3.**
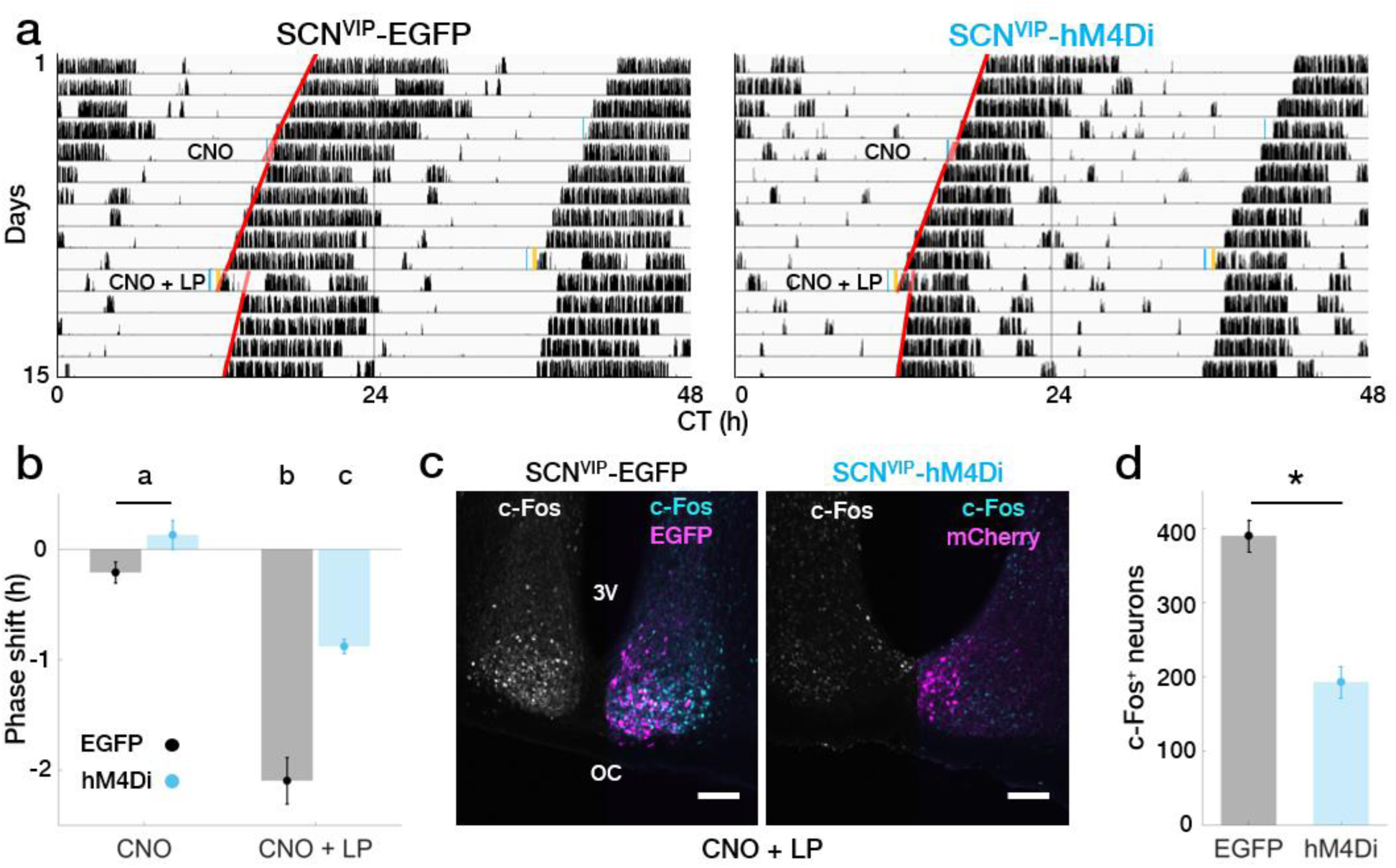
SCN VIP neuron activity is necessary for normal light-induced phase shifts. **a**) Representative double-plotted actograms from transgenic mice free-running in DD without (EGFP) or with (hM4Di) SCN VIP neurons expressing an engineered inhibitory Gi-coupled receptor. Mice were given 1 mg/kg clozapine-n-oxide (CNO) alone or CNO plus a 15-min light pulse (LP) at CT 12. Blue, time of CNO administration; yellow, 15 min bright light pulse; red, activity onset lines of best fit. **b**) CNO attenuated phase delays in response to a light pulse in hM4Di-expressing mice (n = 4 mice per group; Two-Way Repeated Measures ANOVA, F(1, 6) = 28.97, p = 0.0017 with post-hoc Sidak’s MCT). Error bars depict means ± SEM and letters indicate statistically different groups. **c**) Representative confocal images of a SCN slice taken from a EGFP- or hM4Di-expressing mouse one hour after CNO and a 15-min light pulse. Grayscale shows c-Fos expression alone and colored images show c-Fos (cyan) and EGFP or hM4Di-mCherry (magenta) coexpression. 3V, third ventricle; OC, optic chiasm. Scale bar=100 μm. **d**) VIP neuron inhibition attenuated light-induced c-FOS expression (n = 4 mice per group; Student’s t-test, p = 0.0006). Error bars depict means ± SEM.

## DISCUSSION

Together, these experiments demonstrate that SCN VIP neurons exhibit circadian rhythms in both spontaneous and light-evoked neuronal activity *in vivo* and in circadian and NMDA-evoked VIP release *in vitro*. Previous studies have reported circadian rhythms in calcium levels of VIP neurons in the isolated SCN *in vitro* (Enoki et al., 2017), but have differed on whether their firing rate is circadian (Fan et al., 2015; Hermanstyne et al., 2016; Webb et al., 2009). Our results are consistent with light- and circadian-evoked VIP release from the SCN (Shinohara et al., 1995, 1999) which has been not been detectable *in vivo* (Francl et al., 2010). Similarly, our results suggest that physiological levels of NMDA induce VIP release by elevating neuronal activity at times of day when neuronal activity is low but not when neuronal activity is already high, consistent with NMDA’s effects on SCN phase shifts (Ding et al. 1994). High concentrations of NMDA, however, likely reduce VIP release during the subjective day by reducing elevated VIP neuron firing through depolarization block (Meijer et al. 1993). Intriguingly, we also find that the circadian activity of VIP neurons rapidly becomes high and arrhythmic in constant light, consistent with results showing that constant light increases levels of VIP (An et al., 2012), but at apparent odds with the sustained circadian locomotor activity for multiple days in LL. We therefore postulate that aberrant lighting can elevate activity in SCN VIP neurons (some of which project to extra-SCN brain regions; (Abrahamson and Moore, 2001)) without immediate effects on circadian behavior, but with potential effects on behaviors such as mood (LeGates et al., 2012).

Our results indicate that activation of SCN VIP neurons is an essential component of the circadian light transduction pathway. Prior studies showed that nocturnal light upregulates markers of activation in VIP neurons (Kuhlman et al., 2003; Romijn et al., 1996). However, it is unclear if all photic signaling must pass through VIP neurons to entrain rhythms in the SCN and behavior. For example, shifts following optogenetic stimulation of SCN neurons *in vitro* can be blocked by VIP receptor antagonists (Jones et al., 2015), and the acute induction of the core clock gene *Per1*, which is preferentially induced in VIP neurons in response to a light pulse, is required for light-induced behavioral phase shifts (Akiyama et al., 1999; Kuhlman et al., 2003). These results indicate that VIP signaling could be essential for photic entrainment. In contrast, *Vip*-deficient mice can still synchronize to a light cycle (although with an 8-h advanced phase angle of entrainment (Colwell et al., 2003)), and we found that inhibition of VIP neurons only halved light-induced shifts in circadian behavior. It is possible that genetic deletion of VIP early in development allows for compensation or that our chemogenetic manipulations did not fully silence all VIP neurons. However, because retinal ganglion cells make contacts on numerous cell types throughout the SCN (Fernandez et al., 2016) and light induces c-FOS broadly in the SCN (Castel et al., 1997), we propose that VIP neurons are a major, but not a unique, communicator of light information to the SCN. Thus, increases in VIP release by the circadian clock during the day or by environmental light at night synchronize SCN cells to each other and to the environmental light cycle, but the phase of the daily rhythm is likely shaped by additional signals within the SCN.

## METHODS

### Animals

All experiments were performed using roughly equal numbers of male and female mice in accordance with Washington University’s Institutional Animal Care and Use Committee guidelines. No differences in results were observed between sexes. For *in vivo* fiber photometry and chemogenetics experiments, we used heterozygous VIP-Cre knock-in mice (*Vip*^*tm1(cre)Zjh*^/J; Jackson Laboratories 010908; (Taniguchi et al. 2011)) on a C57BL/6JN background that were at least 6 weeks old at the time of virus injection and ~2-5 months old during recordings. To help identify VIP cells in some experiments, mice were crossed to Ai9 or Ai32 (B6.Cg-*Gt(ROSA)26Sor*^*tm9(CAG-tdTomato)Hze*^/J or B6;129S-*Gt(ROSA)26Sor*^*tm32(CAG-COP4*H134R/EYFP)Hze*^/J; Jackson Laboratories 007909 or 012569) mice to yield heterozygous VIP-IRES-Cre; Ai9 or VIP-IRES-Cre; Ai32 mice that expressed a tdTomato or EYFP fluorescent reporter in VIP neurons. For *in vitro* SCN recordings, we collected SCN slices from wild-type or *Vip*^-/-^ (founders generously provided by C. Colwell, University of California, Los Angeles) mice (17-50 days old) on a C57BL/6JN background. After weaning and prior to recording, up to 5 mice of the same sex were housed per cage in a 12:12 light/dark (LD, where lights on is defined as zeitgeber time (ZT) 0; light intensity ~2 × 10^14^ photons/cm^2^/s) cycle with food and water *ad libitum* and constant temperature (~22°C) and humidity (~40%).

### Virus injection and stereotaxic surgery

Cre-inducible AAV vectors encoding GCaMP6s (AAV9-CAG-Flex-GCaMP6s-WPRE-SV40) and EGFP (AAV9-CAG-Flex-EGFP-WPRE-bGH) were obtained from the University of Pennsylvania Vector Core (AV-9-ALL854 and AV-9-PV2818) and used at a final concentration of ~1 × 10^13^ genome copies (GC)/ml. Cre-inducible AAV vectors encoding hM4Di (AAV8-hSyn-DIO-hM4D(Gi)-mCherry) were obtained from Addgene (#44362) and used at a final concentration of ~2 × 10^12^ GC/ml. At the start of surgery, mice received preoperative analgesia (carprofen, 5 mg/kg), were anesthetized with isoflurane (induction dose ~3%, maintenance dose ~1.5%), and placed in a stereotaxic frame (Kopf) on a heating pad to maintain body temperature throughout the procedure. We then infused virus to the SCN (-0.46 mm posterior, ±0.15 lateral, - 5.65 ventral from Bregma) at a rate of 0.05 μl/min through a 30-gauge needle attached to a 2 μl syringe (Neuros, Hamilton). Total volume injected for fiber photometry experiments was 0.3 μl unilaterally (GCaMP6s and EGFP) and, for chemogenetics experiments, was 0.3 or 0.5 μl bilaterally (EGFP or hM4Di, respectively). After infusion, the needle was left in place for at least 8 min before slowly withdrawing. For fiber photometry experiments, we subsequently implanted a fiber optic cannula (6 mm in length, 400 μm diameter core, 0.48 NA, metal ferrule; Doric Lenses) immediately dorsal to the virus injection site. Cannulas were secured with a layer of adhesive cement (C&B Metabond, Parkell) followed by a layer of opaque black dental cement (Ortho-Jet, Lang). After surgery, skin was closed with sutures or surgical glue (Vetbond, 3M) and mice were allowed to recover on a heating pad until ambulatory. Mice were then transferred to their home cages for 2-4 weeks to allow for virus expression.

### *In vivo* fiber photometry recording

After recovery and virus expression, mice were transferred to individual, open-topped, cages equipped with running wheels and bedding maintained in a light-, temperature- and humidity-controlled chamber with food and water *ad libitum*. Lights were set to 12:12 LD, DD or LL (light intensity ~2 × 10^14^ photons/cm^2^/s). Freely moving mice were chronically tethered to a fiber optic cable (400 μm core, 0.48 NA, 1.5 m length; Doric Lenses) connected to the implanted optic fiber by a zirconia mating sleeve (Doric Lenses). Running wheel activity was recorded in 6 min bins using Clocklab software (Actimetrics). After at least 3 d of acclimation to tethering, we began fiber photometry recordings using established methods (Eban-Rothschild et al., 2016). To control for motion artifacts, we excited GCaMP6s with its calcium-dependent excitation wavelength of 470 nm and at its calcium-independent isosbestic excitation wavelength of 405 nm (Lerner et al., 2015). Blue 470 nm and violet 405 nm LED light (M470F3 and M405FP1, Thorlabs) were sinusoidally modulated at 211 and 531 Hz, respectively, using a custom Matlab program (Mathworks) and a multifunction data acquisition device (NI PCI-6110, National Instruments). The excitation lights passed through a fluorescence cube (FMC4, Doric Lenses) containing two separate excitation filters (450-490 nm and 405 nm), reflected off a dichroic mirror, and coupled into the tethered fiber optic cable. Light intensity for each wavelength was set to ~30 μW at the termination of the fiber optic cable. Sinusoidally-modulated GCaMP6s fluorescence was collected through the same fiber optic cable, passed through a emission filter (500-550 nm), and focused through a patch cord onto a femtowatt photoreceiver (Model 2151, Newport). The photoreceiver signal was then sent to two lock-in amplifiers (30 ms time constant, SR830, Stanford Research Systems) that were synchronized to 211 or 531 Hz for demodulation. Voltage signals from the amplifiers were collected at 10 kHz using a custom Matlab program and a multifunction data acquisition device and saved to disk as binary files. For spontaneous activity experiments, hourly 10-min recordings were obtained automatically using a custom Matlab program. For light-evoked activity experiments, hourly recordings were obtained similarly with the addition of a 15 s illumination from the chamber lights delivered at 2 min into the 10-min of data acquisition. Animals were excluded from analysis if both the calcium-dependent and calcium-independent signals simultaneously fluctuated by ~100 mV (an indicator that the fiber was positioned below the SCN), if they did not exhibit circadian locomotor activity or if the fiber or virus was off-target as determined by histology (n = 12 excluded out of 44 recorded mice).

### Fiber photometry data analysis

Data files were opened in Matlab and downsampled to 1 kHz for further analysis as in (Zalocusky et al., 2016). To account for different photobleaching dynamics, extracted calcium-dependent (470 nm) and calcium-independent (405 nm) signals were individually fit with an exponential filter and the fitted signals were subtracted from the raw signals. Signals were then band-pass filtered between 0.016 and 2 Hz (spontaneous recordings) or 0.003 and 2 Hz (evoked recordings) and fit to one another using a linear least-squares fit. Change in fluorescence over baseline fluorescence (ΔF/F) was calculated as (488 nm signal - fitted 405 nm signal) / (fitted 405 nm signal). Final values were adjusted so that a ΔF of 0 corresponded to the second percentile of the signal. ΔF/F values were converted to events by counting instances longer than 1 s when ΔF/F exceeded the median plus two standard deviations of the trace or 1.5% ΔF/F, whichever was greater. We report the events per minute and also the integrated calcium values as defined by integrating the ΔF/F values above 0% over 10 min. For light-evoked recordings, response amplitude was defined as the peak ΔF/F during 2 min following the start of the light pulse divided by the median ΔF/F during the 2-min preceding the light pulse, and response duration was defined as the time from the end of the light pulse until the signal returned to the median ΔF/F of the entire recording.

### VIP sensor cell production

VPAC2R is preferentially coupled to a Gαs-adenylate cyclase-cAMP pathway (Langer, 2012). To generate reporter cells (Muller et al. 2014; Gizowski et al. 2016) that respond to VIP through a Gα_q_-phospholipase C-calcium pathway (Offermanns and Simon, 1995; Zhu et al., 2008), HEK293 cells (ATTC) were sequentially transfected with pcDNA3.1(+)-hVIPR2 (neomycin/kanamycin selection sequence replaced with puromycin sequence) and then pcDNA3.1(+)-hGα_15(16)_ plasmids (cDNA Resource Center). 16 μl FuGENE transfection reagent (Promega) and 1 μg plasmid DNA were diluted in Opti-MEM (Thermo Fisher Scientific) to a final volume of 250 μl. Transfection medium in 2 ml Dulbecco’s modified Eagle’s medium (DMEM) with 5% fetal bovine serum (FBS; Thermo Fisher Scientific) was added to HEK293 cells at 60-80% confluency and removed after 15-20 h. Transfected cells were maintained at 37°C and 5% CO_2_ in CO_2_-buffered DMEM with 10% FBS and 1% penicillin/streptomycin (Thermo Fisher Scientific) for 2 d. On day 3, the antibiotic was replaced with 0.4 μg/ml puromycin (Thermo Fisher Scientific). Cells were selected for 10 d. Single-transfected and selected cells were then transfected with the second plasmid as described. Double-transfected cells were selected with 550 μg/ml G418 sulfate (Thermo Fisher Scientific) for 10 d. Doses of puromycin and G418 were determined through a HEK293 kill curve using a range of 0.38-0.44 μg/ml and 200-800 μg/ml, respectively. Double-transfected and selected cells (HEK293 VIPR2+hG+) were kept in media with puromycin and G418 antibiotics until ready for reporter transfection and recording. To verify successful double-transfection of VIP sensors, cells were immunostained with the primary antibodies rabbit anti-VPAC2 (ab28624, Abcam, 1:500) and mouse anti-Gα15 (sc-393878, Santa Cruz, 1:50) and the secondary antibodies Alexa 488 donkey anti-rabbit and Alexa 594 donkey anti-mouse (711-545-152 and 715-585-150, Jackson Immunoresearch, 1:500). 24 h before recording, 5 × 10^4^ cells were plated on glass-bottom dishes coated with a mixture of poly-D-lysine, laminin, and collagen (Sigma). 3-4 h after plating, we transfected cells with the fluorescent calcium reporter pGP-CMV-NES-jRCaMP1b (Addgene; (Dana et al., 2016). Before recording, the media was changed to recording media containing DMEM supplemented with 100 U/ml penicillin, 100 μg/ml streptomycin, 10 mM HEPES, 0.35 g/L NaHCO3, 4.5 g/L D-glucose, 2% L-glutamine, and 1x B27 cell culture supplement (Thermo Fisher Scientific). To determine the responsivity and specificity of the reporter cells, 1 μl of 1, 10, or 100 nM VIP (Bachem) or 100 nM GRP (Sigma) diluted in recording media was pipetted directly above the plated cells and imaged.

### Organotypic SCN culture and *in vitro* VIP sensor recording

SCN slices were obtained using methods modified from (Abe et al., 2002). Mice were anesthetized with CO_2_ and decapitated. Brains were quickly removed and placed in cold oxygenated Hank’s balanced salt solution (HBSS). 250 μm coronal sections containing the SCN were obtained using a vibratome (OTS-5000, Electron Microscopy Sciences). SCNs were dissected and cultured individually on cell culture membranes (PICM0RG50, Millipore), which were then placed into a 35 mm Petri dish (Falcon) with 1 ml recording medium, sealed with vacuum grease, and incubated at 37°C overnight. The following day, culture membranes containing the SCN were transferred to dishes containing cultured VIP sensor cells. We extrapolated subjective time of day *in vitro* from the prior light cycle. For time-lapse recordings, dishes were placed on an inverted microscope (Eclipse TE2000-S, Nikon) within an incubation chamber (In Vivo Scientific) held at 25°C. VIP sensor fluorescence was imaged with a 10x objective focused on the reporter cells, a TRITC filter cube (Chroma), and a digital EMCCD camera (iXon DU-897, Andor Technology), recorded automatically at 1 frame/sec for 10 min/h for 24 h using a custom Micro-Manager script (http://micro-manager.org). Recordings were stopped after 24 h before reporter cells reached confluence. For acute recordings, VIP sensor cells were prepared as above and kept at 25°C until they were imaged with SCN slices taken at either 11am-1pm (CT6) or 5-7pm (CT 12) the previous day. We imaged VIP sensor fluorescence for 5-8 min using a TRITC filter cube and a digital CCD camera (QIClick, QImaging) at 1 frame/sec with QCapture (QImaging) and CamStudio software. We then pipetted 1 μl of 1 or 10 nM *N*-methyl-D-aspartic acid (NMDA; Sigma) directly onto the SCN slice and recorded for an additional 8-10 min. In all experiments, dishes were excluded from analysis if the VIP sensors did not respond to exogenous VIP application (n = 1 out of 27 dishes).

### VIP sensor analysis

Frames were extracted from movie files using Matlab and opened as TIFF stacks in ImageJ (http://imagej.nih.gov/ij) for analysis. Recordings were bleach corrected using an exponential fit and background subtracted using a rolling ball algorithm to eliminate SCN autofluorescence. Regions of interest were selected using a standard deviation projection and a diameter threshold of 5 to 30 pixels2. Mean pixel intensity values for each ROI calculated for each frame were then processed in Matlab by first dividing by the mean pixel value of the entire frame to generate a background-corrected pixel intensity value. ΔF/F for each ROI for a given frame *n* was then calculated as (corrected pixel intensity value for frame *n* - median pixel intensity values across all frames) / (median pixel intensity values across all frames). Resulting ΔF/F values were smoothed by a 5-s moving average filter. For recordings of spontaneous activity, as there was no pre- or post-stimulus epoch, data were analyzed by integrating the single-cell ΔF/F over each hourly 10-min recording. This single-cell integrated fluorescence was summed and divided by the number of cells analyzed in that hour to account for reporter cell proliferation to produce an average integrated fluorescence per cell. For evoked responses, the maximum ΔF/F for each cell was calculated before and after NMDA stimulation.

## *In vivo* VIP neuron inhibition

After recovery, mice were transferred to wheel-running cages and housed in DD in a light-tight, temperature- and humidity-controlled chamber with food and water provided *ad libitum*. After allowing mice to free-run for multiple days, we injected them with clozapine-N-oxide (CNO, 1 mg/kg; Hello Bio) intraperitoneally 30 min prior to activity onset (Grippo et al., 2017). Mice were returned to their cages and allowed to free-run for multiple days and then injected again with CNO at CT 11.5, 30 min prior to a 15-min light pulse (bright (ranging from 5 × 10^15^ to 2 × 10^16^ photons/s/cm^2^ throughout the cage) or dim (~4 × 10^14^ photons/s/cm^2^) light pulse, returned to their cages and allowed to free run for multiple days. Phase shifts were determined by using activity onset analysis in Clocklab, using at least 4 days of data per condition (baseline, after CNO, and after CNO + light pulse). Days of drug administration were excluded from analysis. A regression line was fit through activity onsets and extrapolated before and after each condition; phase shift duration was determined by calculating the time difference in regression lines extrapolated to the days of drug administration (i.e., the time difference between the predicted and actual phase onsets). To determine light-evoked activation of VIP neurons, mice were injected with CNO at CT 11.5, received a 15-min bright light pulse at CT 12, returned to DD, and perfused 1h after the start of the light pulse as described below.

### Histology and image analysis

Mice were deeply anesthetized with isoflurane (>5%) and transcardially perfused with PBS followed by 4% paraformaldehyde. Brains were extracted, postfixed overnight in 4% paraformaldehyde, and cryoprotected in 30% sucrose. 40 μm coronal sections containing the SCN were collected using a cryostat (Leica) and stored at 4°C in PBS. Immunostaining was performed as previously described (Grippo et al., 2017; Tso et al., 2017). Primary antibodies used were rabbit anti-mCherry (ab168453, Abcam, 1:1000) and goat anti-c-Fos (sc52-G, Santa Cruz, 1:1000). Secondary antibodies used were DyLight 594 donkey anti-rabbit (SA5-10040, Thermo Fisher Scientific; 1:500), DyLight 488 donkey anti-goat (SA5-10086, Thermo Fisher Scientific; 1:500), and CF 647 donkey anti-goat (SAB4600175, Sigma; 1:500). 2048 x 2048 pixel images of the SCN were acquired on a Nikon A1 confocal microscope as a single z-stack (optical section thickness of 2 μm) using identical capture settings for each image that was to be quantitatively compared. For quantitative c-Fos analysis, sections were z-projected by their average intensity using ImageJ software. Regions of interest were selected automatically in ImageJ using Shanbhag thresholding with identical settings for each section followed by particle analysis with size and circularity thresholds of ≥ 200 pixels^2^ and ≥ 0.1, respectively. Brightness and contrast settings were adjusted identically for sections depicted in representative images.

### Statistics

Sample sizes were chosen to be sufficient for statistical analysis based upon similar techniques used in previous publications (e.g. (Eban-Rothschild et al., 2016)). Data analysis was performed blind to genotype or condition. There were no methods to randomize mice to experimental groups or to blind investigators to genotype or condition during the experiments. All statistical analyses including One-Way ANOVA, Two-Way ANOVA, One-Way Repeated Measures ANOVA, Two-Way Repeated Measures ANOVA, Student’s *t*-test, Tukey’s multiple comparison test (MCT), and Sidak’s MCT were performed in Matlab R2016a or Prism 7 (Graphpad) with α defined as 0.05. Circadian rhythmicity was determined using Cosinor analysis (Refinetti et al., 2007) in Matlab and independently using JTK Cycle (Hughes et al., 2010) in R. Shapiro-Wilk and Brown-Forsythe tests were used to test for normality and equal variances. Data are presented as means ± SEM.

## ACKNOWLEDGEMENTS

We thank A. Damato for assistance with chemogenetics experiments and the members of the Herzog lab for discussion and comments on the manuscript. This work is supported by US National Institutes of Health grants R01 NS09536702 (E.D.H), U01 EB02195601 (E.D.H), and F32 HL133772 (J.R.J).

## AUTHOR INFORMATION

### Affiliations

Department of Biology, Washington University in St. Louis, St. Louis, Missouri, USA Jeff R Jones, Tatiana Simon, and Erik D Herzog

Neuroscience Graduate Program, Washington University in St. Louis, St. Louis, Missouri, USA Lorenzo Lones

### Contributions

J.R.J., T.S., L.L., and E.D.H designed the experiments. J.R.J. and E.D.H. analyzed data. J.R.J. and L.L. performed the fiber photometry experiments. J.R.J, T.S., and L.L. performed the VIP sensor experiments. J.R.J. performed the chemogenetics experiments. J.R.J., T.S., L.L., and E.D.H prepared the manuscript.

### Corresponding author

Correspondence to Erik D Herzog

